# Meningococcal Deduced Vaccine Antigen Reactivity (MenDeVAR) Index: a Rapid and Accessible Tool that Exploits Genomic Data in Public Health and Clinical Microbiology Applications

**DOI:** 10.1101/2020.08.18.256834

**Authors:** Charlene M.C. Rodrigues, Keith A. Jolley, Andrew Smith, J. Claire Cameron, Ian M. Feavers, Martin C.J. Maiden

## Abstract

As microbial genomics makes increasingly important contributions to clinical and public health microbiology, the interpretation of whole genome sequence data by non-specialists becomes essential. In the absence of capsule-based vaccines, two protein-based vaccines have been used for the prevention of invasive serogroup B meningococcal disease (IMD), since their licensure in 2013/14. These vaccines have different components and different coverage of meningococcal variants. Hence, decisions regarding which vaccine to use in managing serogroup B IMD outbreaks require information about the index case isolate including: (i) the presence of particular vaccine antigen variants; (ii) the expression of vaccine antigens; and (iii) the likely susceptibility of its antigen variants to antibody-dependent bactericidal killing. To obtain this information requires a multitude of laboratory assays, impractical in real-time clinical settings, where the information is most urgently needed. To facilitate assessment for public health and clinical purposes, we synthesised genomic and experimental data from published sources to develop and implement the ‘Meningococcal Deduced Vaccine Antigen Reactivity’ (MenDeVAR) Index, which is publicly-available on PubMLST (https://pubmlst.org). Using whole genome sequences or individual gene sequences obtained from IMD isolates or clinical specimens, MenDeVAR provides rapid evidence-based information on the presence and possible immunological cross-reactivity of different meningococcal vaccine antigen variants. The MenDeVAR Index enables practitioners who are not genomics specialists to assess the likely reactivity of vaccines for individual cases, outbreak management, or the assessment of public health vaccine programmes. MenDeVAR has been developed in consultation with, but independently of, both vaccine manufacturers.

## Introduction

Microbial whole genome sequencing (WGS) has advanced our understanding of microbial evolution, diversity, and pathogenicity. Since the first bacterial genome was sequenced in 1995, the technology has developed from dideoxynuclueotide terminator (Sanger) sequencing to multiplexed WGS platforms (1-3). Concomitantly, the cost of WGS has reduced substantially, increasing its availability and affordability worldwide; however, DNA sequencing itself is a first step, with multidisciplinary expertise required to exploit these complex large data sets to address particular questions and translate the results to public health action. Genomic technologies are increasingly incorporated into public health and clinical microbiology laboratories, where identifying and typing micro-organisms is critical to informing infectious disease management in individuals and populations. Extracting information from genomic data is important, but it is equally important to communicate these data promptly and effectively to relevant practitioners (4). Here we describe a generalizable framework for assimilating sequence data with phenotypic information, linking genotype to phenotype with the results presented as an easy to understand result for use by non-specialists.

Invasive meningococcal disease (IMD), caused by *Neisseria meningitidis*, is a serious infection with significant mortality and morbidity (5, 6). Diagnosis of IMD is either through bacterial culture and capsular group serotyping, or, in the absence of culture, by PCR testing, with additional discrimination provided by characterisation of capsule-encoding and protein antigen-encoding genes (7). IMD generally occurs sporadically but can occur in clusters and outbreaks, due to the transmission of hyperinvasive meningococcal variants generally or among individuals living in closed- or semi-closed communities such as schools, universities, military barracks, and extended households. Increasingly, real-time WGS of meningococcal isolates can direct public health investigations and interventions.

Prevention of IMD is possible by immunisation, delivered either by routine programmes or in response to clusters or outbreaks. When they occur, such outbreaks are a public health priority, requiring the rapid identification of individuals at high risk from the meningococcal variant identified in the index case. Prophylactic antibiotics are provided to close contacts to prevent outbreak strain transmission and vaccination offered where appropriate (8). While highly immunogenic conjugate protein-polysaccharide vaccines are available against invasive meningococci expressing capsular serogroups A, C, W, and Y (9), there are none against serogroup B meningococci, which are a major cause of IMD outbreaks and clusters in many countries. In 2013 and 2014, two protein-based meningococcal vaccines were licensed to assist in the prevention of serogroup B IMD. The particular protein antigens contained in the two vaccines, 4CMenB (Bexsero®) and rLP2086 (Trumenba®), were different and not specific to serogroup B meningococci. These antigens also displayed immunologically significant protein sequence diversity (10, 11). Therefore, the two vaccines exhibit different degrees of possible protection against heterologous vaccine antigens and, consequently, there could be a need for frontline clinical and public health specialists to assess each vaccine rapidly in the context of a particular scenario to inform decisions about vaccine implementation.

Using WGS to provide clinically applicable information requires systematic and reproducible characterisation of genetic variation. Multilocus sequence typing (MLST), based on housekeeping genes, is the most widely-used approach to characterising bacterial variants, facilitating communication among laboratories internationally and the identification of hyperinvasive meningococci (12). Typing bacterial genetic diversity of medically important features, such as polysaccharide capsules (13, 14), antimicrobial resistance genes (15), and vaccine antigens (16) can be achieved through similar gene-by-gene approaches (17). For example, the Bexsero® Antigen Sequence Typing (BAST) scheme was established to characterise and describe vaccine antigen variants, using data derived through WGS or sequencing of individual genes (16).

Both vaccines contain factor H binding protein (fHbp), one recombinant peptide variant in Bexsero® (peptide 1) and two native lipidated peptide variants in Trumenba® (peptides 45 and 55) (11). Bexsero® also contains the recombinant proteins, Neisserial heparin-binding antigen (NHBA, peptide 2) and *Neisseria* adhesin A (NadA, peptide 8), combined with the PorA-containing (variable region (VR2), peptide 4) outer membrane vesicle from the MeNZB vaccine (10). The BAST scheme catalogues peptide presence/absence and variation, using deduced peptide sequences, but cannot infer protein expression or cross-reactivity. The Meningococcal Antigen Typing System (MATS) laboratory assay was devised to estimate the proportion of diverse serogroup B disease strains prevented by Bexsero®, by assessing protein expression and cross-reactivity (18); however, MATS is not widely or immediately available in clinical settings and is time- and resource-intensive. Genetic MATS (gMATS) was developed to predict Bexsero® strain coverage using sequence and phenotypic MATS data. At the time of writing, this algorithm was not available on an accessible, integrated platform for genome sequence data analysis, nor had it been updated to accommodate the description of additional variants (19).

To perform genomic vaccine antigen analysis comprehensively requires an understanding of sequencing technology, genomic data quality control, and gene/peptide curation and analysis. As of mid-2020, these skills were developing amongst healthcare scientists/clinicians, but were far from universal (4). Given the need to assess breadth of vaccine reactivity and to ensure genomic data are harnessed to maximise clinical and public health benefit, we developed the ‘Meningococcal Deduced Vaccine Antigen Reactivity’ (MenDeVAR) Index, publicly-accessible on PubMLST *Neisseria* website (20). By synthesising published, peer reviewed, experimental data with sequence data, the MenDeVAR Index provides a means for public health and clinical practitioners to extract easily understood, relevant information from genomic data in real-time.

## Methods

### Vaccine antigen typing

Allele-based typing schemes for each of the antigens included in Bexsero® and Trumenba® have been published. The BAST scheme was developed as a multi-locus, rapid, and scalable method to catalogue deduced peptide diversity of meningococcal vaccine antigens (16). The scheme includes five peptide components contained in the Bexsero® vaccine: fHbp; NHBA; NadA; and PorA VR1 and VR2. Typing of Trumenba® vaccine antigen fHbp was available with cross-referencing to the subfamily A and B nomenclature, on PubMLST *Neisseria* website (21, 22). Novel peptide variants are curated in real-time after submission to PubMLST, these curated databases form the basis of the MenDeVAR Index.

### Literature search

Determining the extent to which either protein-based vaccine is protective against a given meningococcus requires an assessment for each vaccine component of the protein sequence variant present, its surface expression, its likely recognition by vaccine-induced antibodies, and finally the likelihood of bactericidal killing of the meningococcus in the presence of vaccinee serum. These factors were assessed using published experimental studies for each vaccine. For Bexsero®, the MATS assay was used, which was established to assess the breadth of vaccine coverage to diverse meningococcal strains (18, 23). MATS determines the antigenic variants of fHbp, NHBA, and NadA through sandwich ELISA, and their reactivity to pooled toddler serum (post-vaccination with three doses and booster), based on a collection of reference strains tested in serum bactericidal activity (SBA) assays. For Trumenba®, the Meningococcal Antigen Surface Expression (MEASURE) assay (24), a flow cytometric measurement of fHbp surface expression, was used. Additionally, SBA assays using serum from individuals immunised with Trumenba® (2 or 3 doses, varying dosing schedules) were included, as there was only one vaccine antigen. Only antigens tested in these assays were analysed as contributing to a cross-protective vaccine effect for the MenDeVAR Index (Figure 1).

**Figure 1:**
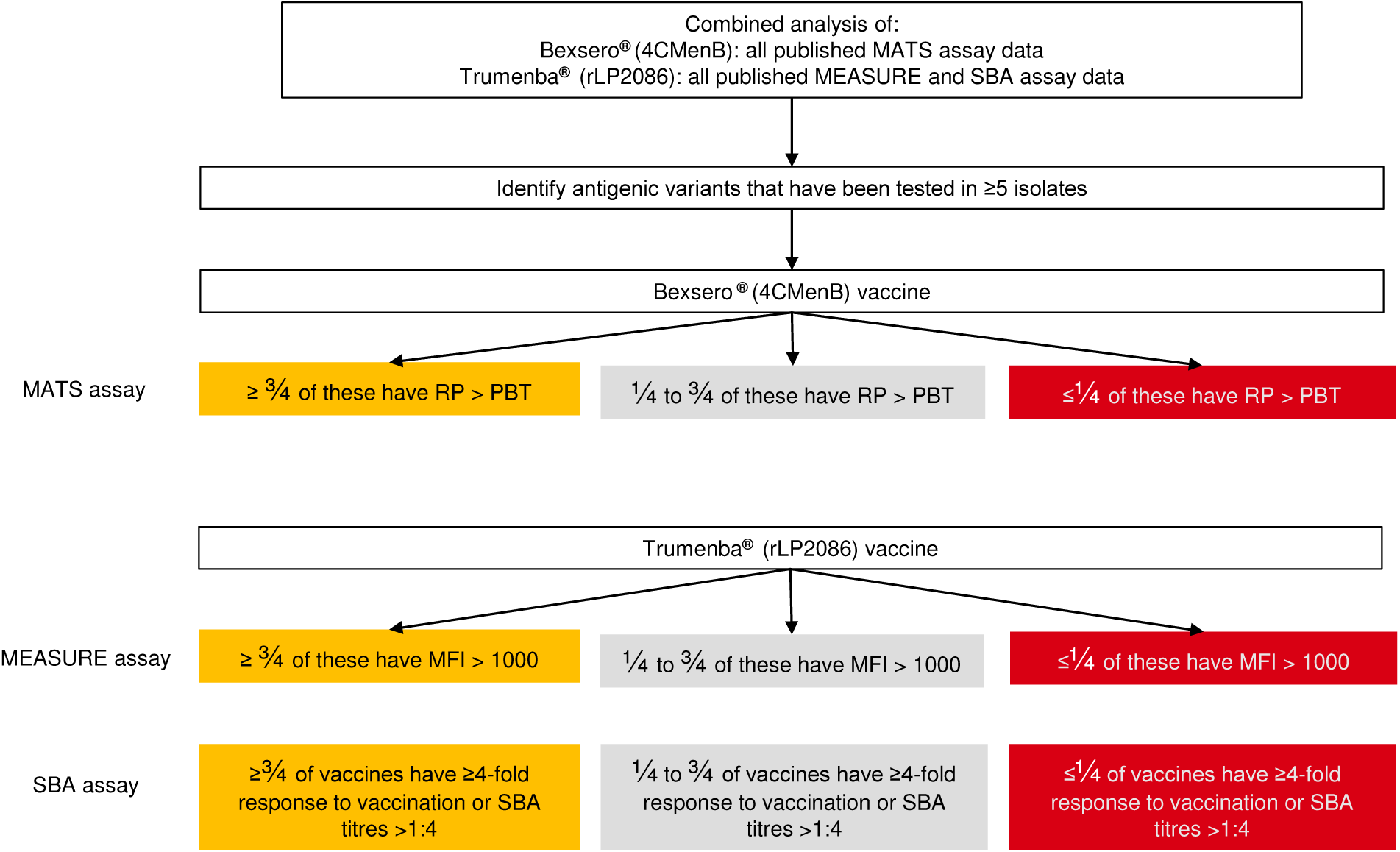
The Meningococcal Deduced Vaccine Antigen Reactivity (MenDeVAR) Index algorithm used to identify which antigens are included as cross-reactive in the combined analysis of published experimental data from: Meningococcal Antigen Typing System (MATS)^18^; Meningococcal Antigen Surface Expression (MEASURE) assay^24^; and serum bactericidal activity (SBA) assay^27^. RP (relative potency), PBT (Positive bactericidal threshold), MFI (mean fluorescence intensity).

For Bexsero®, a literature search using the terms “Meningococcal Antigen Typing System” AND “*Neisseria meningitidis*” AND “vaccine” on 14^th^ May 2020 yielded 44 studies published in English. There were 13 studies eligible for assessment (supplementary Table 4), pertaining to capsular group B IMD isolates (MATS is only validated for serogroup B), with data of sufficient detail to assess individual antigens and their predicted vaccine coverage. For Trumenba®, a literature search using the terms “meningococcal antigen surface expression (MEASURE) assay” AND “*Neisseria meningitidis*” AND “vaccine” on the 14^th^ May 2020 yielded 12 studies published in English. One study contained MEASURE assay data for individual antigenic variants (Table 2). Additionally, a literature search using the terms “serum bactericidal activity assay” AND “*Neisseria meningitidis*” AND “vaccine” AND “bivalent” on the 14^th^ May 2020 yielded 28 studies published in English. Fifteen studies contained data to assess individual antigenic variants and their likelihood of providing protection using SBA assays (supplementary Table 5).

**Table 1:** The caveats that are listed on the PubMLST *Neisseria* website when interpreting the MenDeVAR (Meningococcal Deduced Vaccine Antigen Reactivity) Index. PMID (Pubmed identifier)

|  | Bexsero® vaccine | Trumenba® vaccine |
| --- | --- | --- |
| **Source of data** | These data combine multiple sources of information including: peptide sequence identity through whole genome sequencing; experimental assays developed as indirect measures of the breadth of vaccine protection against diverse meningococci; and assays developed to assess immunogenicity. |  |
| **Protein expression** | We have not inferred protein expression from genomic data, therefore there may be isolates that possess genes but do not express the protein *in vivo*. |  |
| **Cross-reactivity definition** | An antigenic variant was considered cross-reactive if it had been tested in ≥5 isolates/subjects and was above the accepted threshold in ≥3/4 of those isolates. This was established through combined analysis of published experimental studies (PMID provided for each variant), not from genomic data. |  |
| **Meningococcal isolate source** | These assays were based on serogroup B disease isolates for both vaccines. |  |
| **Experimental assays** | • Meningococcal Antigen Typing System (MATS) assay. | • Meningococcal antigen surface expression (MEASURE) assay.<br>• Serum bactericidal activity (SBA) assay. |
| **Age of vaccinees** | • For MATS assay development, Bexsero® vaccine recipients were infants who had received 3 doses of vaccine and then a booster at 12 months.<br>• The pooled sera used for the MATS assay were taken from the toddlers at 13 months of age. | • The age of vaccine recipients in the experimental studies varies widely, ranging from toddlers to adults, and needs to be taken into consideration when interpreting results.<br>• Vaccine studies used different schedules and doses of vaccines. |

**Table 2:** Vaccine antigen variants for the protein-based meningococcal vaccines Bexsero® (4CMenB) and Trumenba® (rLP2086) and their designation by Meningococcal Deduced Vaccine Antigen Reactivity (MenDeVAR) Index as: “green”, exact matches to the sequence variants; “amber”, cross-reactive in experimental studies; “red”, not cross-reactive in experimental studies; “grey”, insufficient data”. fHbp, factor H binding protein; NHBA, Neisserial heparin-binding antigen; NadA, Neisseria adhesin A; PorA VR2, Porin A variable region.

| MenDeVAR Index | Antigen requirement | fHbp | NHBA | NadA | PorA VR2 |
| --- | --- | --- | --- | --- | --- |
| **Bexsero® vaccine** |  |  |  |  |  |
| **Green**<br>**(exact)** | ≥1 of: | 1 | 2 | 8 | 4 |
| **Amber**<br>**(cross-reactive)** | ≥1 of: | 4, 10, 12, 14, 15, 37, 110, 144, 215, 232 | 1, 5, 10, 113, 243, 607 | 3, 6 | - |
| **Red**<br>**(none)** | All 4: | 16, 19, 21, 22, 24, 25, 29, 30, 31, 45, 47, 59, 76, 109, 119 | 6, 9, 17, 18, 25, 30, 31, 43, 47, 63, 112, 120, 160, 187, 197 | 1, 21, 100 | ≠4 |
| **Grey**<br>**(insufficient data)** | None of the above | 13, 321 and other antigens that have not been experimentally tested | 3, 20, 21, 24, 29, 115, 118, 130 and other antigens that have not been experimentally tested | Any antigens that have not been experimentally tested | - |
| **Trumenba® vaccine** |  |  |  |  |  |
| **Green**<br>**(exact)** | 1 of: | 45, 55 |  |  |  |
| **Amber**<br>**(cross-reactive)** | 1 of: | 1, 4, 13, 14, 15, 16, 19, 21, 23, 25, 30, 47, 49, 76, 87, 180, 187, 252, 276, 510 |  |  |  |
| **Red**<br>**(none)** | 1 of: | - |  |  |  |
| **Grey**<br>**(insufficient data)** | None of the above | 13, 24 or other antigens that have not been experimentally tested |  |  |  |

### Criteria for defining cross-reactive antigens in the MenDeVAR Index

To index the experimental data, thresholds were determined to define antigenic variants as either likely cross-reactive or not cross-reactive and the proportion of isolates with a given antigenic variant considered covered/protected through experimental assays was calculated. For each assay (MATS, MEASURE, and SBA), thresholds previously defined by the developers or the research community were employed. For the MATS assay, an antigenic variant was considered “covered” (i.e. would be susceptible to a vaccine-induced immune response) where the relative potency (RP) was greater than the positive bactericidal threshold (PBT) (18). For the MEASURE assay, an antigenic variant was considered “covered” if the mean fluorescent intensity (MFI) >1000 (24). For the SBA assay, antigenic variants were assessed through host immunogenicity, resulting in likely protection from infection. The accepted serological measure indicating likely protection by immunisation with meningococcal vaccines is either ≥4-fold rise in antibody titres between pre- and post-vaccination sera or a titre >1:4 (27, 28). From the combined analysis of the experimental studies, if an antigenic variant had been tested in ≥5 isolates and ≥3/4 of them were covered/protected, then the variant was considered cross-reactive (“amber”). If an antigenic variant had been tested in ≥5 isolates, and ≥3/4 of them were not covered/protected, then the variant was considered not cross-reactive (“red”) (Figure 1).

### Development of data visualisation for the MenDeVAR Index

For ease of data presentation, a red, amber, green ‘traffic light’ data interpretation was employed: “green” was assigned to meningococcal variants with ≥1 antigenic vaccine variant, based on exact peptide sequence match; “amber” was assigned to isolates with ≥1 antigenic variant demonstrated as cross-reactive in experimental studies; and, “red” was assigned to isolates where all antigenic variants were not exact matches and had been shown to not elicit cross-reactivity to vaccine variants. The designation “grey” was assigned to variants possessing antigenic variants untested in experimental assays at the time of writing or where such tests did not meet the threshold chosen to indicate cross-reactivity. The MenDeVAR Index status of the variants, especially those designated as “grey”, will be updated in the light of the above criteria as further published information become available.

The MenDeVAR Index was implemented on the PubMLST *Neisseria* website on the isolate record (Figure 2a). In addition, WGS data or individual gene sequences can be used to make a direct query on https://pubmlst.org/bigsdb?db=pubmlst_neisseria_mendevar, which outputs the MenDeVAR Index result, without the need to create isolate records or upload WGS data to the database (Figure 2b). A written description is provided to aid those with colour vision deficits, where “green” means “exact”, “amber” means “cross-reactive”, “red” means “none”, and grey means “insufficient data”. Additional, supporting information is provided: (i) the antigenic determinant of the reactivity index; (ii) the assay used to determine cross-reactivity; (iii) specific references to studies including those antigens; and (iv) caveats to interpretation (Table 1).

**Figure 2:**
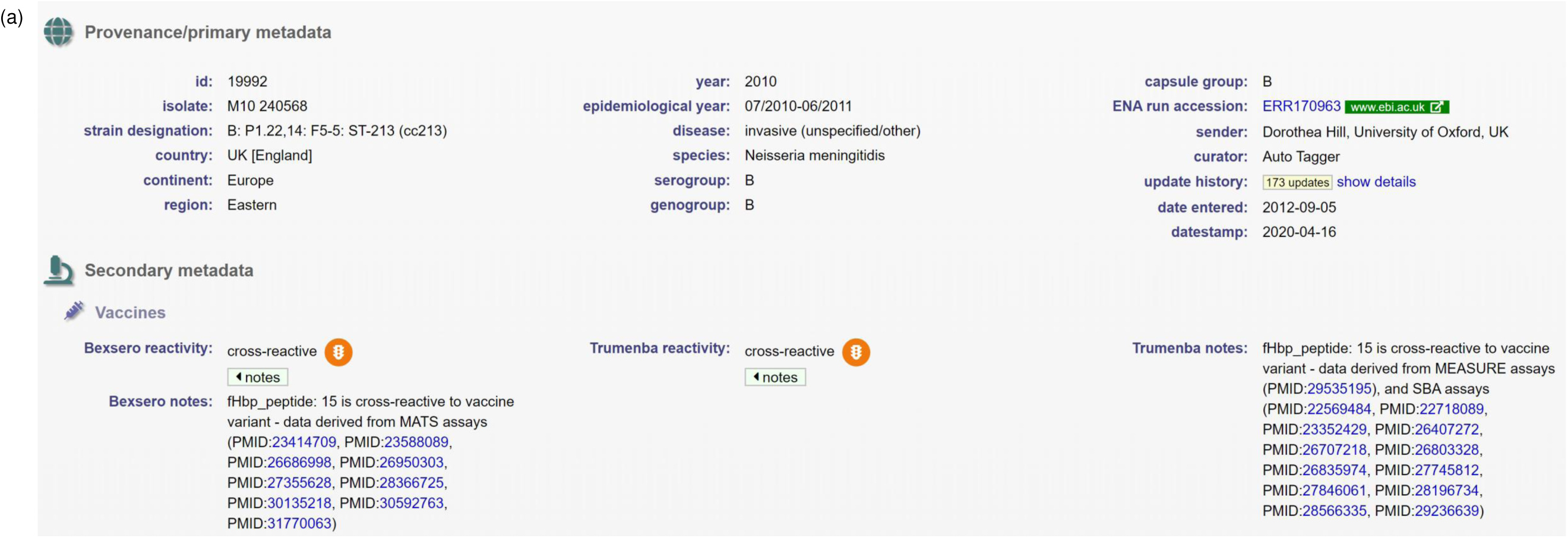
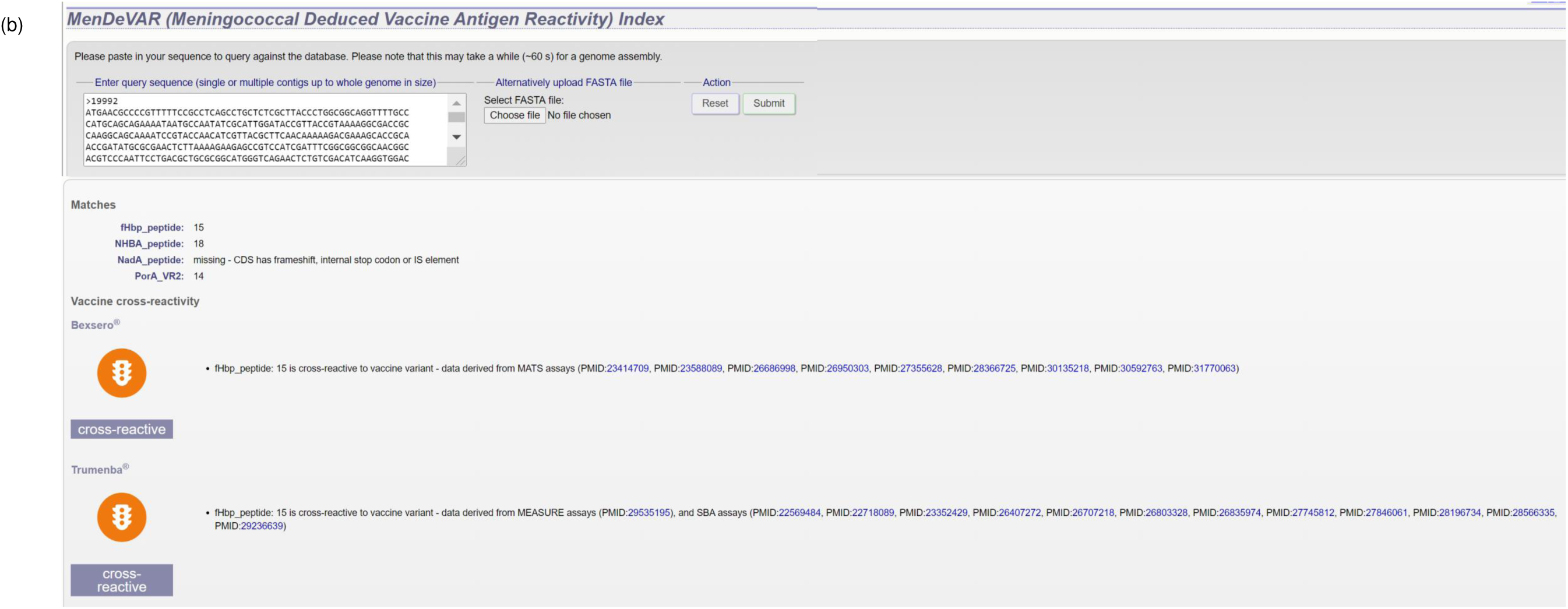
(a) The Meningococcal Deduced Vaccine Antigen Reactivity (MenDeVAR) Index as it appears on the isolate record page of the https://PubMLST/org/neisseria website. The provenance data shows the PubMLST id is 19992, states this is a serogroup B meningococcal isolate from the Eastern region of the UK, collected from invasive disease in 2010. The MenDeVAR Index is shown under the secondary metadata heading, and shows this isolate contains cross-reactive antigens for both vaccines, with fHbp peptide 15 the antigen used to determine this through the MATS assay for Bexsero® and the MEASURE and SBA assays for Trumenba ®, reference shown with PubMed ID (PMID). (b) The web interface to search using genome sequence, individual genes or whole genome data to output the MenDeVAR Index.

### Case studies

To exemplify the application of the MenDeVAR Index, two published IMD outbreaks/clusters were analysed: a IMD outbreak amongst a semi-closed, Irish traveller community (2010-2013) (25); and a university IMD outbreak in the USA (2016) (26). Both WGS data available through PubMLST and published antigenic variants determined through WGS were examined.

## Results

### Cross-reactive vaccine antigens

For Bexsero® vaccine, MATS studies (29-41) were identified through literature searches. With the exception of two studies (34, 40) that used PBT for fHbp of 0.012, all other antigen RP were assessed against the PBT of 0.021 for fHbp, 0.294 for NHBA, and 0.009 for NadA (18). For each antigenic variant of fHbp, NHBA, and NadA, the proportion of isolates with a RP>PBT was calculated. For fHbp, there were 139 peptides examined by MATS assay, 28 (20.1%) tested in ≥5 isolates. For NHBA there were 110 peptides, 30 (27.3%) tested in ≥5 isolates. For NadA, there were 22 peptides, 5 (22.7%) tested in ≥5 isolates. For Trumenba® vaccine, each antigen tested by the MEASURE assay in one study (24) was evaluated. For fHbp, there were 9 peptides examined by MEASURE assay, 6 of which were tested in ≥5 isolates (Table 3). From SBA studies (42-56), there were 23 fHbp peptides examined by SBA assay, 23 (100.0%) tested in ≥5 isolates.

**Table 3:** Two examples of outbreak/clusters from published literature, showing the molecular typing data used to determine the MenDeVAR (Meningococcal Deduced Vaccine Antigen Reactivity) Index. ST, sequence type; cc, clonal complex; PorA VR, Porin A variable region; FetA, enterobactin receptor FetA; fHbp, factor H binding protein; NHBA, Neisserial heparin-binding antigen; NadA, *Neisseria* adhesin A; BAST, Bexsero® Antigen Sequence Type.

| Cases | Year | Capsular group | ST | cc | PorA typing | FetA typing | fHbp peptide | NHBA peptide | NadA peptide | PorA VR1 | PorA VR2 | BAST | Bexsero® MenDeVAR Index | Trumenba® MenDeVAR Index | PubMLST id |
| --- | --- | --- | --- | --- | --- | --- | --- | --- | --- | --- | --- | --- | --- | --- | --- |
| Irish traveller community outbreak |  |  |  |  |  |  |  |  |  |  |  |  |  |  |  |
| Case A | 2010 | B | Incomplete MLST profile | ST-41/44 complex | 7-2,4 | F1-21 | no data | no data | no data | 7-2 | 4 | - | Green | Grey | - |
| Case B | 2010 | B | 6697 | ST-41/44 complex | 7-2,4 | F1-21 | 4 | 607 | 0 | 7-2 | 4 | 381 | Green | Amber | 26834 |
| Case C | 2011 | B | 6697 | ST-41/44 complex | 7-2,4 | F5-12 | no data | no data | no data | 7-2 | 4 | - | Green | Grey | - |
| Case D | 2012 | B | Incomplete MLST profile | - | - | no data | no data | no data | no data | no data | no data | - | - | - | - |
| Case E | 2013 | B | 6697 | ST-41/44 complex | 7-2,4 | F5-12 | no data | no data | no data | 7-2 | 4 | - | Green | Grey | - |
| Case F | 2013 | B | Incomplete MLST profile | ST-41/44 complex | 7-2,4 | no data | no data | no data | no data | 7-2 | 4 | - | Green | Grey | - |
| Case G | 2013 | B | No data | - | - | no data | no data | no data | no data | no data | no data | - | - | - | - |
| Case H | 2013 | B | 6697 | ST-41/44 complex | 7-2,4 | F5-12 | 4 | truncated | 0 | 7-2 | 4 | Incomplete BAST profile | Green | Amber | 30743 |
| US university cluster |  |  |  |  |  |  |  |  |  |  |  |  |  |  |  |
| Case 1 | 2016 | B | 11 | 11 | 5-1 , 10-1 | no data | 19 | 20 | 0 | 5-1 | 10-1 | 3545 | Grey | Amber | - |
| Case 2 | 2016 | B | 11 | 11 | 5-1 , 10-1 | no data | 19 | 20 | 0 | 5-1 | 10-1 | 3545 | Grey | Amber | - |

Antigenic variants that did not meet either cross-reactive or not cross-reactive threshold were designated as “grey”, indicating that insufficient data were available to make an assessment for this variant. This included variants: (i) tested in ≥5 isolates, with between 1/4 and 3/4 covered/protected (Table 2); (ii) tested in <5 isolates (for Bexsero® vaccine this was 111 fHbp peptides, 80 NHBA peptides, and 17 NadA peptides, and for Trumenba® vaccine 3 fHbp peptides tested by MEASURE assay); or (iii) not tested in experimental assays.

### Designation of isolates with the MenDeVAR Index

A meningococcal variant was designated “green” if it contained at ≥1 exact sequence match to the vaccine antigenic variants. This was, for Bexsero®: fHbp peptide 1; NHBA peptide 2; NadA peptide 8; and PorA VR2,4 (16, 57). Similarly, for Trumenba® this corresponded to fHbp peptides 45 or 55 (11) (Table 2). The “amber” designation was a used if a meningococcus contained ≥1 antigenic variant deemed cross-reactive from experimental studies, from any of fHbp, NHBA or NadA (Table 3). PorA peptides are not considered cross-reactive (58). Finally, the “red” designation was used for meningococci where none of its antigens present were exact matches with the vaccine antigens and its antigen variants had been shown experimentally not to cross-react with antibodies elicited by the vaccine (Table 2).

### MenDeVAR Index: exemplar case studies

#### Irish traveller community outbreak

Retrospective analysis of a published IMD outbreak in the Republic of Ireland (2010-2013) (25), exemplified the potential use of the MenDeVAR Index in the context of a community outbreak, where a variety of clinical specimens were available. A total of eight cases were identified over 42-months (Table 4). The initial meningococcus, from Case A, was not cultured, but identification and typing data were acquired by PCR amplification and sequencing of MLST loci and fine-typing antigen-encoding genes *porA* and *fetA*. PorA VR2 antigen 4 was present, an exact peptide sequence match to Bexsero®. There was insufficient data to inform the use of Trumenba®, which contains only fHbp proteins. At the time of identification Case A was considered to be sporadic case and the appropriate public health action was antibiotic prophylaxis for close contacts. Using the MenDeVAR Index, the disease-associated meningococcus would have be designated “green” for Bexsero® and “grey” for Trumenba®. Of the seven cases subsequently linked to this case, only two were successfully cultured and WGS (Cases B and H), but five could have a MenDeVAR Index inferred from fine-typing antigen PorA, with respect to Bexsero® (Table 4). Additional molecular fHbp typing of isolates would inform the use of Trumenba®, in a setting where the PorA is not variant 4. These data identified 75% (6/8) of isolates, two with WGS, with sufficient information to designate as MenDeVAR Index “green” for Bexsero® and two WGS isolates with “amber” for Trumenba® (Table 3).

#### US university cluster

A cluster of IMD occurring in the US (2016) (26) was examined to demonstrate the use the MenDeVAR Index in an institutional outbreak. In this cluster, two undergraduate students at a New Jersey university were diagnosed with serogroup B IMD, with meningococci isolated from the cerebrospinal fluid of both (26). These isolates were examined in real-time by WGS through the local public health department and were both sequence type 11 (clonal complex 11) and indistinguishable (Table 3). Antigenic variant data provided in the publication was assessed, which provided data equivalent to that obtained by determining the antigenic variants through PCR and sequencing, if WGS had not been available. The meningococci causing the outbreak harboured fHbp variant 2 peptide 19, an antigen which is cross-reactive with Trumenba® (“amber”) but not cross-reactive with Bexsero® (“red”). The outbreak strains also had: (i) no *nadA* gene present (“red”); (ii) PorA 10-1 (“red”); and (iii) NHBA peptide 20, for which there is insufficient data to determine cross-reactivity with confidence (“grey”). The MenDeVAR Index therefore designated these isolates “amber” for Trumenba® and “grey” for Bexsero®, the latter based solely on the NHBA variant present, with remaining antigens “red”. This information could have directed public health specialists to using Trumenba® early after IMD cluster definition was met, preventing delays in health protection interventions including mass vaccination campaigns, frequently required in university settings.

## Discussion

As bacterial genome sequencing has become increasingly accessible, the prospect of using genomic data for the benefit of public and individual health has become a reality. This opportunity is, however, fraught with challenges including: (i) the large and complex genomic datasets involved; (ii) the expertise required to understand the uses and limitations of WGS technologies; (iii) the increasing number and complexity of analysis tools; (iv) the requirement for skills with command-line interfaces; (v) insufficient bioinformatics or genomic epidemiology training amongst healthcare practitioners and scientists; and (vi) the diversity of the information sources that need to be integrated.

Genome sequence data provide information on the presence or absence of genes associated with clinically relevant phenotypes e.g. antibiotic susceptibility, pathogenicity or vaccine antigens. The first step in exploiting this information is to extract relevant data for the identification of the genes and the protein variants they encode (typing). The second step is to index these types to the relevant phenotypic data. The third step is to present the result in an accessible format for non-genomics specialists to inform clinical decision-making. Here, we demonstrated the MenDeVAR Index, which combines these steps into a system for rapid, real-time assessment of protein-based meningococcal vaccine antigens, for public health and clinical microbiology application.

The epidemiology of IMD varies geographically. Sporadic cases occur in countries where IMD is endemic, with clusters and outbreaks associated with high-density living conditions such as universities, military, or travelling communities (59). Endemic and hyper-endemic serogroup B IMD is problematic in many industrialised regions (60) and, in the absence of group B polysaccharide vaccines, protein-based vaccines (10, 11) have been developed. When IMD outbreaks emerge, it is essential to identify contacts and implement public health interventions rapidly. These include antibiotics and vaccinations, the latter, especially, requiring timely serogroup determination of the outbreak strain to ensure deployment of the appropriate vaccine (8). For serogroup B outbreaks, characterisation of peptide antigens is required to assess whether vaccination with Bexsero® and/or Trumenba® is likely to prevent disease (8). At the time of writing, this assessment was only possible using the laboratory assays established during the clinical development of these vaccines to assess their breadth of antigenic coverage, namely the MATS, MEASURE, and SBA assays (18, 24, 28). These assays, however, required growth of the causative isolate, were confined to reference laboratories in a limited number of countries, and were time-consuming and expensive to perform (26, 61). Consequently, they could not be relied upon to inform timely public health interventions. At the same time, WGS has become increasingly accessible to microbiology laboratories, often in real-time or near real-time. Further, where meningococcal cultures were not available, PCR of fine-typing and fHbp antigens provided information that complements the phenotypic data compiled within the MenDeVAR Index. Interpreted by local microbiologists and epidemiologists in the context of other pertinent information, the MenDeVAR Index offers a pragmatic assessment of likely susceptibility of outbreak strains to vaccine-induced immunity, based on published data.

For the development of the MenDeVAR Index, robust, pragmatic criteria were used to assess the weight of evidence of potential antigenic cross-reactivity from four different sources. The SBA titre remained the accepted immune correlate of protection for assessing meningococcal vaccine efficacy; however, the SBA assay cannot be performed for routine IMD case isolates investigated as part of a public health response for many reasons including the availability of expertise, resources, time, human complement, and infant sera. The use of MATS and MEASURE assays, as means of assessing the breadth of antigenic coverage, generated the best data available. Data from MEASURE assays, however, were limited at the time of assessment and the MATS assay was suggested to provide a conservative estimate compared to SBA titre (36, 38, 41). SBA data were not included for Bexsero®, which as a multi-component vaccine could induce multiple antibody responses. Although the gMATS assay also used genotypic predictors of MATS phenotype, and predicted cross-reactivity in agreement with the MenDeVAR Index using similar criteria (fHbp peptides: 1, 4, 10, 12, 14, 15, 37, 110, 144, 215, 224, 232; NHBA peptides 1, 5, 10, 113, 243, 607) (19), the gMATs system was only applicable to one of the two available protein-based vaccines. Moreover, it excluded NadA antigens as predictors, included some unpublished data, and had not been updated. The MenDeVAR Index can assist public health and microbiology specialists by compiling and indexing the complex data available in the published evaluation of hundreds of meningococcal antigenic variants, a total of 29 studies at the time of writing.

The MenDeVAR Index is accessible through a user-friendly webpage (https://pubmlst.org/bigsdb?db=pubmlst_neisseria_mendevar) that facilitates the submission of WGS data as single or multiple contigs, or as part of an isolate record on PubMLST *Neisseria* website.

The case studies explored here demonstrated how the MenDeVAR Index can be used as outbreaks developed, with the Irish outbreak showing how multiple types of information can be used effectively. Had the MenDeVAR Index been available at the time, it would have supported the use of the Bexsero® vaccine in this outbreak setting. The US university cluster demonstrated the difficulties faced by public health specialists in combining complex datasets from multiple sources in real-time to inform intervention strategies. This cluster was investigated by US Centers for Disease Control and the isolates were sent for laboratory testing at US universities, which is not a routine procedure. These analyses identified relatively low fHbp protein expression and low binding of NHBA peptide 2 antisera to the outbreak strain, suggesting reduced likelihood of bactericidal killing (26). Based on these data along with additional information about persistence of antibody responses post-vaccination, immunisation of ∼35,000 university students with Trumenba® was recommended. The public health team acknowledged that WGS data indicated the presence or absence of particular antigenic variants, which could be compared to the respective vaccine antigens. When variants were not exact sequence matches, however, there was no additional information available to indicate potential cross-protection offered by the vaccine. In the case of this outbreak, the MenDeVAR Index would have supported the use of Trumenba®, solely on the basis of WGS data.

There are limitations to using the MenDeVAR Index, as it is based on WGS data linked to information from published *in vitro* MATS, MEASURE, and SBA serological studies, (Figure 3). These assays are not perfect surrogates of protection for a variety of reasons including the age groups used to establish the assays and the provenance of the isolates used in their development. Further, at the time of writing, the expression of the antigens could not be reliably inferred or predicted from WGS data, although some fHbp promoter and intergenic regions had been correlated with protein expression (65, 66). Finally, the MenDeVAR Index applies to only to possible direct protection against IMD, with no information available about possible herd immunity due to the lack of evidence to suggest either vaccine impacted oropharyngeal carriage of serogroup B meningococci (62-64).

In conclusion, we present a generalizable multi-locus gene-by-gene framework for interpreting complex genomic datasets that can be used by practitioners to address clinical questions in a timely manner. Specifically, the MenDeVAR Index combines genomic and experimental data to provide a rational, evidence-based, estimate of the likelihood that either of the meningococcal protein-based vaccines offers protection against a given meningococcus. To ensure broad accessibility, the MenDeVAR Index is implemented with a ‘red’, ‘amber’, and ‘green’ interpretive interface that is easy to use and informative for practitioners without expertise in genomic analysis. In the light of new published evidence, the MenDeVAR Index can be regularly re-evaluated using the criteria described here, adjusting antigenic variant designations accordingly, to ensure that public health and clinical microbiologists globally benefit from the latest research findings.

## Acknowledgements

This publication made use of the Neisseria Multi Locus Sequence Typing website (https://pubmlst.org/neisseria/) sited at the University of Oxford (Jolley *et al*. Wellcome Open Res 2018, 3:124. The development of this site has been funded by the Wellcome Trust and European Union.

## Supplementary data

**Table 4:** Experimental studies identified through literature search to determine the cross-reactive antigenic variants to Bexsero® (4CMenB) vaccine included for combined analysis.

**Table 5:** Experimental studies identified through literature search to determine the cross-reactive antigenic variants to Trumenba® (rLP2086) vaccine included for combined analysis.

